# Protocol-Guided Cross-Domain Transfer Learning for Bovine Facial Pain Recognition under Weak Dairy-Farm Labels

**DOI:** 10.64898/2026.06.18.733162

**Authors:** Shivam Patel, Suresh Neethirajan

## Abstract

Livestock welfare models are developed under controlled experimental conditions but deployed across farms, breeds, management systems and label regimes, where reliability remains uncertain. We introduce the Protocol-Driven Transfer Evaluation (PDTE) framework, which treats the adaptation protocol, comprising label mapping, objective design, domain alignment, model selection, calibration and threshold policy, as the experimental variable and evaluates transfer through animal-level external validation with uncertainty quantification. We apply PDTE to a bovine welfare task involving transfer of a facial pain representation from postoperative beef cattle to dairy cows under shifts in breed, sex, production system, clinical etiology, recording environment and label fidelity. Using an author-collected Canadian Holstein and Jersey dataset with an independent eight-cow test cohort, direct source-domain transfer was weak, with sequence AUC 0.418 and cow-level AUC 0.400. PDTE identified two failure modes under weak supervision: threshold collapse, in which adaptation converges to a single prediction class, and calibration-induced collapse, in which score ranking is preserved while decision behavior deteriorates. Across protocols, objective design dominated performance. Class-balanced focal adaptation achieved stable operating behavior (sequence AUC 0.611; cow-level AUC 0.667), while a target-only model attained comparable performance without source initialization (sequence AUC 0.596; paired p = 0.984), indicating that protocol design and operating-point choices contributed more than pretraining under weak-label conditions. Animal-level uncertainty remained substantial, with a bootstrap 95% confidence interval of 0.20 to 1.00, exceeding the transfer effect. These findings show that transferability limits cannot be inferred from source-domain performance alone and require protocol-controlled, uncertainty-aware evaluation in livestock AI.

## 1. Introduction

Pain compromises both the welfare and productivity of dairy cattle, yet reliable on-farm detection remains difficult. As prey animals, cattle often suppress overt pain behaviours, and routine assessment still depends on observer-scored instruments whose interpretation varies among raters and contexts (Tschoner et al., 2024; Fischer-Tenhagen et al., 2022). Common dairy conditions, particularly lameness and mastitis, alter behaviour and pain sensitivity and can benefit from analgesia (Flower et al., 2008; Peters et al., 2015). Timely, objective and scalable detection is therefore a meaningful welfare, productivity and management goal (Roche et al., 2025).

Automated facial-video analysis offers a promising route to such detection, and deep learning has made it increasingly feasible to recover pain-related signals from animal video (Broome et al., 2023; Chiavaccini et al., 2024). In cattle, video-based models have performed competitively with trained veterinarians when labels, acquisition conditions and evaluation criteria are tightly controlled (Feighelstein et al., 2026). The central barrier is generalisation. Models developed in controlled studies are eventually expected to operate across different breeds, facilities, disease states, camera conditions and label regimes. Whether a learned representation remains informative after this move, rather than merely performing well within its original dataset, is the question that determines practical value.

This problem is especially acute for bovine pain assessment. The most rigorously annotated cattle pain datasets come from controlled postoperative studies using validated scales such as the UNESP-Botucatu Cattle Pain Scale (UCAPS) (de Oliveira et al., 2014; Tomacheuski et al., 2023). Commercial dairy-farm datasets, in contrast, usually contain heterogeneous recording environments, variable image quality, multiple disease aetiologies and weak health-status labels rather than time-aligned pain scores. Moving from postoperative beef cattle to dairy-farm welfare monitoring therefore introduces simultaneous domain shifts in breed morphology, sex, production role, clinical context, environmental conditions and label fidelity. Transfer-learning theory predicts that such shifts can erode source-domain performance unless adaptation explicitly addresses source-target mismatch (Pan and Yang, 2010; Ben-David et al., 2010; Wang et al., 2022). Similar degradation has been observed in human facial-pain recognition and other computer-vision settings when models are evaluated outside their original data distributions (Othman et al., 2019; Yu et al., 2022).

Despite growing interest in livestock computer vision, little is known about how facial pain representations behave when moved across independent bovine populations, production systems and clinical contexts. Most studies emphasise within-dataset performance, whereas deployment requires reliability under biological and operational variability. More importantly, previous work rarely separates the contribution of the representation from the contribution of the adaptation strategy. Apparent transfer gains may reflect reusable source-domain knowledge, but they may also arise from objective design, threshold selection, calibration, model selection or other protocol choices. Distinguishing these possibilities is essential before transfer claims can be treated as evidence of robust livestock welfare AI.

Addressing this gap requires moving beyond conventional benchmark evaluation toward systematic characterisation of transfer behaviour under realistic agricultural conditions. In practical livestock systems, dense expert pain annotations are rarely available; weak health-event labels and small animal cohorts are often the only supervisory substrate. The central question is therefore not whether a large, ideal, fully annotated dairy-cattle dataset could support accurate pain recognition. It is whether source-domain knowledge, adaptation design and decision policy can be disentangled when limited weak labels are all that farms can realistically provide.

To address this problem, we introduce the Protocol-Driven Transfer Evaluation (PDTE) framework. Existing animal-welfare transfer studies often report final performance without separating the protocol choices that shape transfer behavior, making it difficult to determine whether an apparent gain reflects reusable source-domain knowledge, weak target supervision, or artifacts of objective design, calibration, thresholding, or model selection. PDTE holds the network architecture fixed and treats the adaptation protocol as the experimental instrument, decomposing transfer into six controllable components: label mapping (M), training objective (L), domain-alignment term (A), model-selection rule (S), calibration (C), and operating-threshold policy (T). Each configuration is then evaluated under strict animal-level external validation with cohort-level uncertainty. We apply PDTE to a deliberately demanding UCAPS-to-dairy transfer case: a fixed spatiotemporal facial model initialized from a validated postoperative beef-cattle pain task is adapted to an author-collected Canadian dairy-farm dataset of Holstein and Jersey cows, including healthy animals and animals affected by lameness or mastitis, under weak health-status labels. The aim is not to claim that source pretraining solves dairy-cow pain recognition, but to determine whether apparent transfer survives protocol-controlled, uncertainty-aware evaluation. Accordingly, the target task is framed conservatively as recognition of pain-related discomfort signs, and all target-domain results are interpreted as proxy discrimination rather than validated pain detection.

PDTE differs from reporting a single end-to-end result in three ways that the field did not previously have in one place. First, it decomposes adaptation choices into explicit, auditable components (M, L, A, S, C, T) so that differences in outcome can be attributed to specific decisions instead of unstructured implementation changes. Second, it builds animal-level, uncertainty-aware external validation into the design, so that transfer claims are evaluated at the level of cows with explicit confidence intervals rather than only sequence-level scores. Third, it surfaces two failure modes: threshold collapse and calibration-induced collapse that remain invisible if one reports AUC or accuracy alone. Respectable ranking performance can coexist with operationally degenerate farm-monitoring behavior when threshold selection and calibration are not evaluated jointly with discrimination; PDTE makes that distinction measurable.

The contributions of this work are fourfold. First, we provide one of the first systematic evaluations of bovine facial-pain representation transfer across simultaneous shifts in breed, production system, clinical aetiology, recording environment and label quality. Second, we introduce PDTE as a protocol-guided framework that decomposes adaptation into auditable, reproducible components, enabling investigators to determine whether apparent transfer gains are genuine, weak or protocol-driven. Third, we identify characteristic failure modes in weak-label livestock welfare AI, including threshold collapse, calibration-induced collapse and decision instability despite preserved ranking. Fourth, by including animal-level uncertainty, paired protocol comparisons and a target-only baseline, we isolate the relative roles of source-domain pretraining and protocol design in determining observed performance.

Collectively, this work advances precision livestock computing by providing a reusable framework for evaluating and stress-testing animal-welfare AI systems under weak supervision and agricultural domain shift. The study is deliberately demanding: an eight-cow held-out dairy cohort labelled only by weak health-status proxies, a setting representative of real farms but insufficient for deployment claims. Its value is therefore not a claim that transfer learning has solved dairy-cow pain recognition, but a principled demonstration of how transferability, decision reliability and animal-level uncertainty should be examined before welfare AI systems are translated to farms. The remainder of the paper reviews related work (Section 2), describes the source and target materials (Section 3) and the PDTE methodology and evaluation (Section 4), reports the external-validation results (Section 5), and discusses what PDTE reveals about failure modes, uncertainty, and the limits of cross-domain bovine facial pain transfer for precision livestock welfare AI, with limitations and conclusions (Sections 6-8).

## 2. Related Work

### 2.1. Pain assessment and facial expression in cattle

Reliable labels are the foundation of any transfer study, yet pain labels in cattle remain difficult to obtain. Most reference instruments depend on human observation, and systematic reviews report variability across raters, scales and contexts (Tschoner et al., 2024). Facial-expression scoring is attractive because it is non-invasive, but its reliability varies across large domestic animals, and grimace-scale performance depends on species, pain type, observer training and setting (Fischer-Tenhagen et al., 2022; Evangelista et al., 2022). Validated cattle pain scales perform well in controlled postoperative settings (Tomacheuski et al., 2023), which is the source-domain regime used in this study. Dairy-farm pain, however, is more heterogeneous, arising from lameness, mastitis, systemic inflammation and chronic discomfort rather than a single controlled stimulus (Roche et al., 2025; Ginger et al., 2023; Linstadt et al., 2024). Facial changes during inflammatory mastitis are measurable but entangled with posture, behaviour and clinical state (Ginger et al., 2023). Work on calf grimace scales and broader domestic-animal pain scales further shows that facial signals can overlap with stress and other negative affective states (Farghal et al., 2024; Mota-Rojas et al., 2025). This source-target asymmetry, validated postoperative pain labels versus weak farm health-state proxies, defines the measurement challenge addressed here.

### 2.2. Vision-based recognition of animal pain

Computer vision has expanded animal-welfare assessment from tracking and behaviour recognition toward inference of internal states such as pain. Surveys highlight the promise of these methods but also recurring constraints: small datasets, subtle visual cues, sensitivity to recording conditions and limited evidence of cross-context generalisation (Broome et al., 2023; Chiavaccini et al., 2024). Strong in-domain results have been reported in rabbits, cats and goats, and broader pain-AI work has explored multimodal fusion of facial video with physiological signals (Feighelstein et al., 2023; Martvel et al., 2024; Chiavaccini et al., 2024b; Gkikas et al., 2024; El-Tallawy et al., 2024). In cattle, deep video models have been compared with trained veterinarians under controlled protocols (Feighelstein et al., 2026), facial changes have been quantified during induced mastitis (Ginger et al., 2023), accelerometry has been combined with observation during systemic inflammation (Ledoux et al., 2023), and dairy-cattle micro-expression analysis has been piloted (Zhang et al., 2025). These studies establish that pain-related information can be extracted from animal video. They provide less evidence on whether the same representations remain reliable when herd, breed, recording environment or label definition changes, which is the central deployment question for precision livestock farming.

### 2.3. Transferring pain models across domains, states and morphologies

Evidence on pain-model transfer is stronger in human facial-pain analysis than in livestock. Human studies show that high-performing pain classifiers can lose accuracy across datasets or subject domains, and reviews emphasise subject-independent evaluation, external validation and standardised reporting (Morsali and Ghaffari, 2025; Huo et al., 2024). In animals, transfer evidence is narrower. Cross-state transfer has been shown in horses, where models trained on acute experimental pain helped recognise subtler orthopaedic pain (Broome et al., 2019). Cross-environment studies in sheep show that naturalistic recording conditions degrade pain-recognition performance even with expert annotation (Feighelstein et al., 2025). Weakly supervised domain adaptation has also been explored in human pain estimation by adapting spatiotemporal models from labelled sources to coarse target labels, a setting close to farm deployment (Praveen et al., 2020; Rajasekhar et al., 2021). Cross-morphology transfer remains least developed; horse-to-donkey facial-pain transfer showed that failures can propagate through pose estimation, landmark detection and downstream prediction (Hummel et al., 2020). For cattle, adjacent work supports individual components of the problem, including dairy facial analysis, cattle video pain recognition and micro-expression analysis (Ginger et al., 2023; Feighelstein et al., 2026; Zhang et al., 2025). However, cross-breed, cross-etiology, weak-label transfer to an independent dairy herd remains largely unevaluated.

### 2.4. Transfer learning and domain shift

Transfer-learning theory explains why pretraining alone is not evidence of transferability. Surveys distinguish feature reuse, fine-tuning, domain adaptation and domain generalisation, and show that success depends on source-target relatedness, target-set size and the type of distribution shift (Zhuang et al., 2021; Wang et al., 2022). Medical imaging provides a useful parallel because it also operates with small target cohorts and acquisition heterogeneity; gains depend on task similarity, target-data volume and whether source features are reusable (Kim et al., 2022; Kora et al., 2022; Matsoukas et al., 2022). In bovine facial pain, these issues map directly onto agricultural shifts: imaging and posture changes resemble covariate shift; disease-frequency differences resemble label-distribution shift; and differences in how pain is expressed across pathology or breed approach concept shift. Domain-adaptation theory formalises this by linking target performance to source-target discrepancy and compatibility of labelling functions (Ben-David et al., 2010). Adversarial alignment and correlation alignment are representative strategies, but domain-adaptation reviews caution that alignment helps only when its assumptions hold (Guan and Liu, 2022; Ganin et al., 2016; Sun and Saenko, 2016). External-validation studies in clinical imaging, acquisition-shift correction and cross-species veterinary transfer all show that apparently strong internal performance can fail on independent data (Yu et al., 2022; Roschewitz et al., 2023; Groendahl et al., 2023). The recurring lesson is clear: credible transfer claims require explicit evaluation protocols, not source-domain performance alone.

### 2.5. Research gap and contribution of the present study

Across veterinary pain assessment, animal computer vision and transfer learning, one gap remains unresolved: there is no established framework for testing whether livestock welfare representations transfer under realistic agricultural domain shift. Veterinary pain research clarifies what labels mean but rarely tests whether learned representations survive a change of herd. Transfer-learning theory identifies the shifts that threaten generalisation but is seldom instantiated in weakly labelled farm welfare data with small animal cohorts. Precision-livestock computer vision is advancing rapidly in behaviour recognition, skeleton-based dairy-cow action analysis and video-based lameness detection (Rohan et al., 2024; Hua et al., 2023; Sohan et al., 2026; Tun et al., 2026), yet most systems are evaluated in-domain and focus on visible behaviour, pose or lesions rather than cross-domain reuse of welfare representations.

This study addresses that gap by introducing the Protocol-Driven Transfer Evaluation (PDTE) framework. Rather than proposing another bovine pain classifier, PDTE fixes the architecture and treats the adaptation protocol as the experimental instrument. It evaluates transfer under animal-level external validation, cow-clustered uncertainty, operating-point failure analysis, paired comparisons and a target-only baseline that isolates the contribution of source pretraining. We demonstrate PDTE on a deliberately difficult case: transferring a facial pain representation from postoperative beef cattle to weakly labelled dairy cows under simultaneous biological, environmental and label-quality shifts. The intended contribution is methodological and practical: a reusable way to determine whether apparent welfare-AI transfer is genuine, fragile or protocol-driven before such systems are deployed on farms.

## 3. Materials

### 3.1. Study material overview

This study uses two bovine facial-video domains that differ in population, recording context, pain construct, and label quality. The source domain is a previously published postoperative beef-cattle pain setting associated with the UNESP-Botucatu Cattle Pain Scale (UCAPS), in which Angus and Nelore bulls were assessed around surgical orchiectomy using validated bovine pain instruments (de Oliveira et al., 2014; Tomacheuski et al., 2023). The target domain is an author-collected dairy-cow dataset from the Ruminant Animal Centre (RAC), Dalhousie University, Truro, Nova Scotia, Canada, where health status provides weak supervision for pain-related discomfort rather than time-aligned pain scores. The target population comprises female Holstein and Jersey dairy cows, whereas the source population comprises male beef bulls; sex was not treated as an independent experimental variable but constitutes an additional domain-shift factor in the transfer setting. The contrast between these domains is the empirical basis for the transfer problem: the study does not ask whether pain can be recognized within one carefully controlled dataset, but whether a source facial representation remains informative under dairy-farm morphology, etiology, and label-quality shifts (Figure 1). The attributes that distinguish the source and target domains are summarized in Table 1.

**Figure 1.**
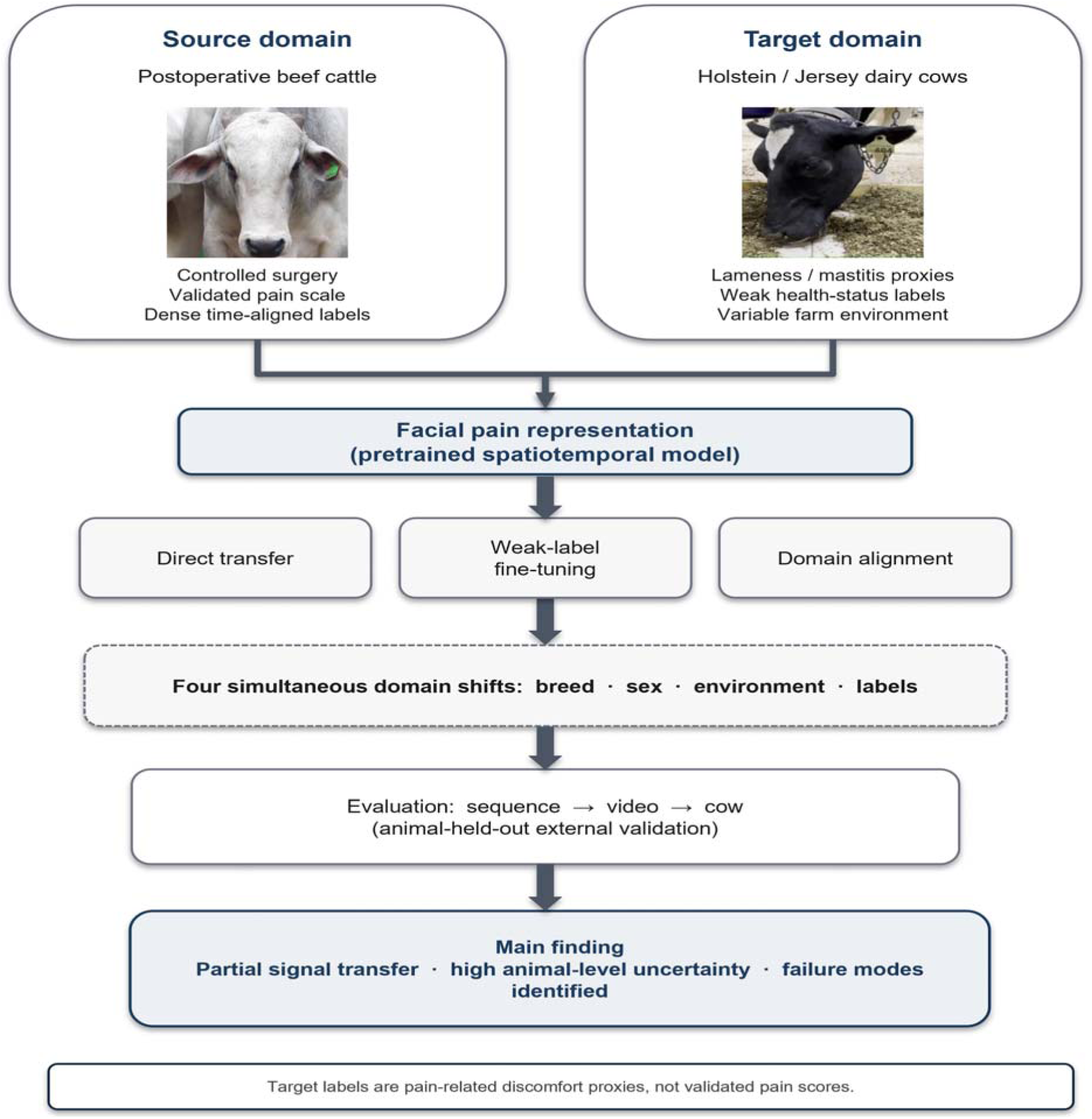
Cross-domain transfer of bovine facial pain representations. A facial pain model pretrained on postoperative beef cattle is transferred to a weakly labeled dairy-cow domain. Transferability is assessed through protocol-guided adaptation and animal-level external validation.

**Table 1.** Source and target domain characteristics for cross-domain transfer learning. Comparison of the beef-cattle source domain and dairy-cow target domain, illustrating the major sources of biological, environmental, and label-related distribution shift.

| Domain attribute | Source domain | Target domain |
| --- | --- | --- |
| Study role | Representation source | Transfer/adaptation and evaluation domain |
| Provenance | Published UCAPS postoperative bull material from UNESP- | Author-collected RAC dairy-cow video at Dalhousie |
|  | Botucatu-associated studies (de Oliveira et al., 2014; Tomacheuski et al., 2023; Feighelstein et al., 2026) | University, Truro, Canada |
| Animal population | Angus and Nelore beef-cattle bulls | Holstein and Jersey dairy cows |
| Primary clinical context | Controlled postoperative pain following surgical castration | Routine dairy-farm health states, including lameness and mastitis |
| Label basis | Validated bovine pain-scale context | Veterinary health-status proxy |
| Label interpretation | Pain-scale-supervised facial pain representation | Pain-related discomfort proxy |
| Recording context | Comparatively controlled facial video | Naturalistic farm video with variation in viewpoint, lighting, and background |
| Main shift axes | Reference domain | Breed/sex, etiology, environment, and label-quality shift |

### 3.2. Source domain: UCAPS postoperative beef-cattle material

The source domain is a controlled postoperative beef-cattle facial pain setting derived from UCAPS-associated studies of bulls undergoing surgical orchiectomy. Its pain construct is grounded in validated cattle pain-scale work, including the original UNESP-Botucatu scale validation and subsequent reliability/validity testing in Bos taurus (Angus) and Bos indicus (Nelore) bulls (de Oliveira et al., 2014; Tomacheuski et al., 2023). Recent automated cattle-pain work has also used UCAPS as the reference standard for comparing deep learning video-based models with trained veterinary assessment (Feighelstein et al., 2026).

For the present Materials section, the relevant source-domain characteristics are its controlled postoperative setting, validated pain-scale supervision, and beef-bull population. These features define the contrast with the target domain, where labels are dairy-farm health-status proxies rather than UCAPS pain scores.

### 3.3. Target domain: RAC dairy-farm video dataset

The target material consists of facial videos of dairy cows collected by the authors at the Ruminant Animal Centre, Dalhousie University, Truro, Nova Scotia, Canada. The recorded population includes Holstein and Jersey cows observed under routine management, exercise, and health-assessment conditions rather than under a controlled surgical protocol. The cohort includes clinically healthy cows and cows identified as affected by disease-associated conditions relevant to pain and discomfort, principally lameness and mastitis, both of which are established painful conditions in dairy cattle (Flower et al., 2008; Peters et al., 2015). The resulting target domain is therefore intentionally realistic: it includes variation in viewpoint, lighting, background, individual morphology, session context, and health condition that would be expected in dairy-farm deployment.

The raw target-video inventory comprised 624 readable videos from 33 parsed cow identifiers. After removing videos without verifiable healthy/unhealthy status and excluding the cow with unknown health status, the classifiable inventory contained 565 videos from 32 cows, with 250 healthy-status and 315 unhealthy-status videos (Table 2). Health status was assigned from recording-session labels when these were explicit and from a provided cow-health mapping when the recording hierarchy encoded date and cow identity rather than health condition.

**Table 2.** Scale and composition of the target video inventory underpinning animal-level external validation, before sequence extraction.

| Inventory level | Value reported in this study |
| --- | --- |
| Collection site | Ruminant Animal Centre, Dalhousie University,<br>Truro, Nova Scotia, Canada |
| Raw readable target videos | 624 |
| Parsed cow identifiers in raw inventory | 33 |
| Classifiable target videos after health-label<br>filtering | 565 |
| Classifiable cows after excluding unknown-health<br>cow | 32 |
| Healthy-status videos in classifiable inventory | 250 |
| Unhealthy-status videos in classifiable inventory | 315 |
| Healthy-status cows in classifiable inventory | 15 |
| Unhealthy-status cows in classifiable inventory | 17 |

Because cows could appear across multiple recording contexts, cow-level health status and sequence-level health status are intentionally distinguished. For cow-level summaries and animal-level partitioning, a cow was treated as unhealthy if it appeared in any documented unhealthy context. For sequence-level supervision, however, each sequence retained the health status of its originating target video. This convention preserves animal-level independence while acknowledging that health state can vary by recording context; its implications are treated as part of the weak-label limitation rather than as clinical ground truth.

For the curated transfer dataset, a fixed selection procedure chose up to three videos per cow before sequence extraction. This cap reduced over-representation of frequently recorded animals while retaining broad cow-level coverage for animal-held-out evaluation. Healthy cows could contribute videos from any available session, whereas unhealthy cows were restricted to during-and after-exercise recordings, with an additional severe-unhealthy event subset included for the affected cow. Cows with fewer eligible recordings contributed all available eligible videos and were retained as partial selections. This procedure selected 92 target videos from 32 cows before sequence-level quality filtering.

The selected videos were converted into short face-centered sequences using sliding temporal windows. The 10 s duration was chosen to retain short facial and postural behavior while remaining compact enough for sequence-level modeling; the 8 s stride produced 2 s overlap between adjacent windows, increasing temporal coverage without making neighboring sequences nearly identical. Each retained window contains 240 stored frames at 24 frames/s. Face detection and quality-control filtering were then applied, retaining only windows that met detection-rate, detection-confidence, and sequence-completeness criteria. The final curated target set contains 549 quality-controlled sequences from 31 cows (Table 3). For model input, each retained sequence is represented by 32 uniformly sampled face crops at 112 x 112 RGB resolution, matching the fixed input format of the source model. The full curation pathway from raw inventory to the held-out cow test set is summarized in Supplementary Figure S1.

**Table 3.** Curation rigor of the target sequence set and the held-out cow material that enables strict animal-level external validation.

| Sequence or split element | Value reported in this study |
| --- | --- |
| Selected target videos before sequence quality control | 92 |
| Selected cows before sequence quality control | 32 |
| Curated target sequences | 549 |
| Cows represented in curated set | 31 |
| Sequence duration | 10 s |
| Frame rate | 24 frames/s |
| Stored frames per sequence | 240 |
| Sequence extraction stride | 8 s |
| Healthy-status proxy sequences | 234 |
| Unhealthy-status proxy sequences | 315 |
| Fixed held-out test cows | 8 |
| Held-out test sequences | 143 |
| Held-out healthy-status proxy sequences | 98 |
| Held-out unhealthy-status proxy sequences | 45 |
| Cross-validation scheme for remaining cows | Five animal-level folds, with four validation cows per fold |
| Model input representation | Uniformly sampled face crops at 32 frames per sequence and 112 x 112 RGB resolution |

The resulting material is imbalanced at more than one level: the curated set contains more unhealthy-status than healthy-status sequences, while the held-out test set is balanced by cow count but not by sequence count. The fixed held-out test set comprises eight cows, with four cows from each health-status branch and 143 sequences in total. The remaining cows are used for animal-level cross-validation and model selection, as described in Section 4. Because the held-out test set contains a modest number of animals, cow-level estimates are interpreted with uncertainty intervals rather than as definitive population-level performance benchmarks.

### 3.4. Weak label semantics and target task

The target labels are weak health-status proxies rather than time-aligned pain scores. Each sequence inherits the health status assigned to its originating target video. Healthy-status sequences are mapped to the no-pain-related-discomfort proxy class, and unhealthy-status sequences are mapped to the pain-related-discomfort proxy class. This binary mapping is intentionally conservative because disease-associated dairy conditions may reflect overlapping pain, discomfort, sickness behavior, stress, and handling context rather than a single isolated pain construct. For this reason, the target task is framed as recognition of pain-related discomfort signs associated with dairy health status. All target-domain results are interpreted as proxy-label discrimination: a positive result indicates information correlated with dairy health-state proxies, whereas a weak or degenerate result indicates source-target mismatch under weak supervision.

### 3.5. Animal-use and ethics approval

Target-domain dairy-cow material. Target-domain facial video was collected at the Ruminant Animal Centre (RAC), Dalhousie University, Truro, Nova Scotia, Canada. All data collection procedures were reviewed and approved by the Dalhousie University Animal Care and Use Committee (Protocol 2024-026; approval date, 16 May 2024) in accordance with Canadian Council on Animal Care (CCAC) guidelines. Data acquisition was entirely non-invasive, involving passive image and video capture without physical contact or intervention with the animals. Participating farm owners provided informed written consent after being fully briefed on the study objectives and protocols. This adherence to ethical standards ensured that animal welfare was not compromised throughout data collection and subsequent computational analysis. Source-domain UCAPS beef-cattle material. The source UCAPS-associated postoperative beef-cattle pain material derives from prior clinical prospective studies approved by the School of Veterinary Medicine and Animal Science, São Paulo State University (UNESP), Ethical Committee for the Use of Animals in Research (approval number 0147/2018). Human ethics approval was not applicable to the source studies. The bulls were part of an approved experiment investigating perioperative pain after surgical castration and were used to validate bovine pain scales. Respecting the three Rs (reduce, replace, and refine), the present study reused existing perioperative pain-assessment observations and video recordings from those approved studies and did not conduct additional source-domain animal experimentation for the transfer experiments reported here.

## 4. Methods

### 4.1. The Protocol-Driven Transfer Evaluation (PDTE) framework

Prior livestock-welfare transfer studies report a single end-to-end performance number, which conflates the many design choices that determine whether a representation actually transfers. We instead adopt the Protocol-Driven Transfer Evaluation (PDTE) framework, which makes those choices the unit of analysis. PDTE holds the network topology and source initialization fixed and treats the adaptation protocol as the experimental instrument, so that every measured difference is attributable to a named, controlled substitution rather than to architecture search or unstructured implementation changes. The decisions PDTE exposes are the six that govern how an available bovine facial-pain representation is reused in a small, weakly labeled dairy-farm target domain: the target-label mapping, loss function, optional domain-alignment term, model-selection rule, calibration procedure, and threshold policy. The fixed spatiotemporal architecture and these variable protocol components are shown in Figure 2.

**Figure 2.**
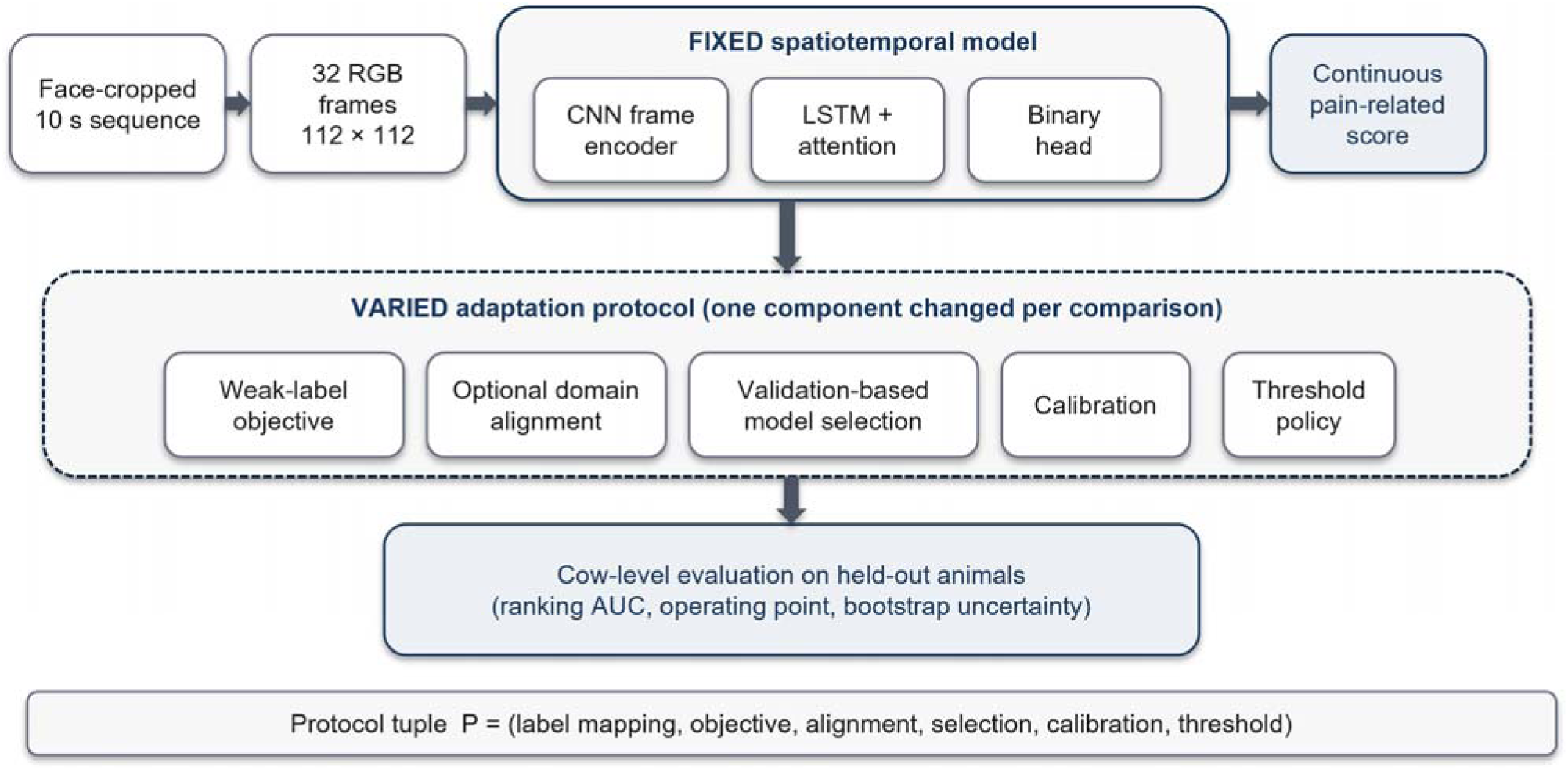
Fixed spatiotemporal architecture and variable transfer protocol. Transfer pipeline used across protocol configurations. Each 10 s face sequence is represented by 32 RGB frames resized to 112 x 112 pixels and passed through a fixed spatiotemporal model consisting of a convolutional frame encoder, recurrent temporal integration, attention pooling, and a binary output score. The evaluated protocol varies the weak-label objective, optional domain-alignment term, validation-selection rule, calibration rule, and threshold policy while keeping the architecture fixed.

### 4.2. PDTE protocol decomposition

PDTE describes each transfer system by a protocol tuple of six adaptation components applied to the same source-initialized model and the same animal-level partitions: the target-label mapping (M), the target objective (L), an optional source-target alignment term (A), the validation-based model-selection rule (S), an optional calibration step (C), and the operating-threshold policy (T). Two systems that share the architecture and data partitions but differ in any one component are treated as distinct transfer protocols, so every result in Section 5 traces to a defined adaptation substitution. This decomposition is the methodological contribution: by separating M, L, A, S, C, and T and evaluating each under identical animal-level external validation, PDTE turns “does the model transfer?” into the more useful question of which adaptation decision governs transfer, and it is what makes the failure modes in Section 5 attributable to specific components rather than to the system as a whole. The formal notation for the protocol tuple and the full per-component specification are given in the Supplementary Methods (Supplementary Table S2). The components instantiated here are the ones relevant to this target domain; PDTE is extensible to additional components as agricultural transfer settings demand.

### 4.3. Base spatiotemporal model and source initialization

The base model consumes a face-centered video sequence and produces one binary logit for the pain-related-discomfort proxy. Each target sequence is represented by 32 uniformly sampled RGB frames resized to 112 x 112 pixels. A two-dimensional convolutional frame encoder maps each frame to a 256-dimensional embedding; a single-layer recurrent module with 128 hidden units integrates the frame embeddings over time; an attention-pooling layer forms a sequence descriptor; and a binary classification head produces the target logit.

All transfer-domain configurations were initialized from a source model trained with the same topology on the UCAPS-associated postoperative beef-cattle pain task (de Oliveira et al., 2014; Tomacheuski et al., 2023; Feighelstein et al., 2026). Source training used animal-held-out folds, 32-frame inputs at 112 x 112 resolution, a binary pain head, and a multi-class source pain-state head; the source model was then reused for target-domain transfer. During transfer adaptation, the convolutional frame encoder was frozen unless otherwise stated, and the temporal, attention, and classification-head parameters were updated. Freezing low-level features is a common small-data transfer practice intended to preserve source-learned visual structure while limiting overfitting on weak target labels (Kim et al., 2022; Kora et al., 2022). In this study, however, the source and target populations differ markedly in coat pattern, head shape, ear position, and facial proportions, so freezing early layers may also prevent adaptation to morphology-specific cues. The target-only baseline in Section 5.8 therefore unfroze the encoder because random initialization requires learning low-level features from the target data; that contrast is methodologically deliberate rather than an inconsistency across experiments.

### 4.4. Transfer-protocol families

Three protocol families were evaluated, together with a calibration analysis; their fixed and varied elements are summarized in Table 4. Direct transfer applied the source model to the target sequences without target-domain parameter updates. Source weights were not optimized on the dairy data, and target labels were not used to fit model parameters. To avoid the substrate mismatch that can arise when source-only baselines are evaluated on earlier target cohorts, the direct-transfer baseline was evaluated on the same curated target sequence set and the same held-out cow test set used for the adaptation protocols. The primary direct-transfer endpoint was the continuous source-model pain-related score, evaluated by AUC on the held-out cows. When thresholded direct-transfer metrics are reported, the threshold is derived only from non-test target animals using the validation policy described below. Direct transfer therefore tests whether the source representation alone ranks dairy health-status proxies above chance and provides the source-only reference condition for the descriptive protocol comparison.

**Table 4.**
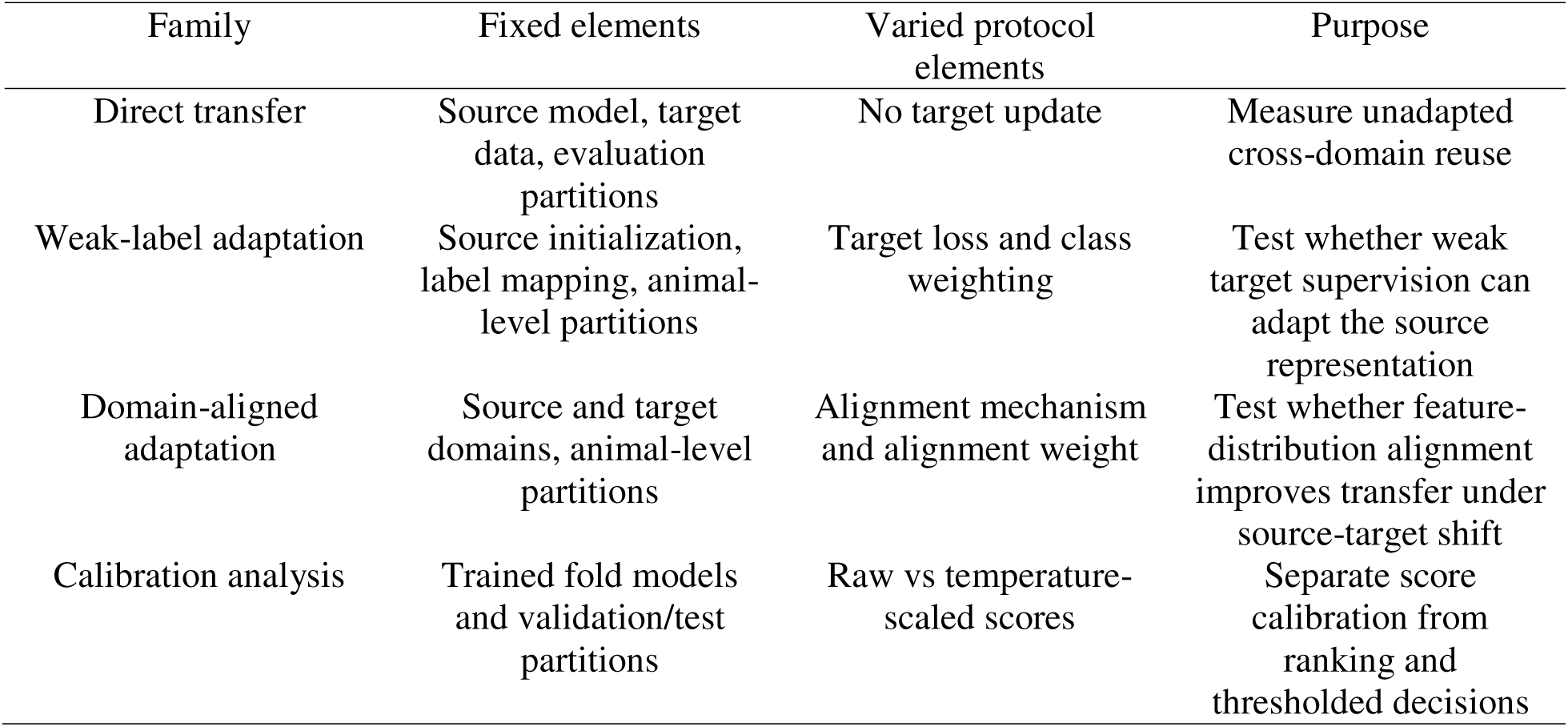
The decomposed PDTE protocol space: controlled substitutions across label mapping, objective, alignment, selection, calibration, and threshold that serve as the methodological instrument of this study.

Weak-label adaptation updated the trainable target-side parameters using the binary health-status proxy labels. The label mapping was kept fixed across weak-label configurations, while the objective varied across binary cross-entropy, focal loss (Lin et al., 2017), and generalized cross-entropy (Zhang and Sabuncu, 2018), each evaluated with and without class-balanced weighting (Cui et al., 2019). The objective definitions and the specific focusing, robustness, and class-balancing hyperparameters are listed in the Supplementary Methods (Supplementary Table S3).

Domain-aligned adaptation added an alignment term A intended to reduce source-target feature-distribution mismatch. Two alignment mechanisms were evaluated: adversarial domain alignment, implemented with a domain classifier and gradient reversal (Ganin et al., 2016), and correlation alignment, which penalizes differences in second-order feature statistics (Sun and Saenko, 2016). Alignment terms were evaluated as protocol components rather than assumed remedies, because domain adaptation can help only when the source and target labeling functions are sufficiently compatible (Ben-David et al., 2010; Guan and Liu, 2022).

### 4.5. Model selection, ensembling, and calibration

All model selection was performed using validation animals only. The selection rule combined three validation quantities, sequence-level AUC, cow-level AUC, and balanced accuracy at an operating threshold, into a single weighted score; the exact weights are given in the Supplementary Methods. The weights are heuristic and were not tuned on the held-out test cows. They give substantial weight to cow-level ranking because animal-level inference is the welfare-relevant unit, retain sequence-level ranking as a secondary signal, and include balanced accuracy to discourage selection of a model with a collapsed operating point. Fold models were selected independently using this validation criterion. Because no formal sensitivity analysis over alternative weights was performed, the resulting performance should be interpreted as the behavior of this prespecified selection rule, not as proof that these weights are optimal or that the same configuration would be selected under all reasonable weighting schemes.

For final held-out inference, fold-level logits were averaged before applying the sigmoid transformation. Averaging logits rather than probabilities keeps the ensemble on the model-score scale and reduces dependence on any single validation split. Temperature scaling was fitted on validation logits for each fold and applied only as a secondary calibrated analysis (Guo et al., 2017). Because temperature scaling is monotonic, it does not change AUC; calibrated and raw operating-point metrics are therefore reported separately.

### 4.6. Threshold policy

The operating threshold T was selected from pooled validation predictions, not from the held-out test set. Candidate thresholds were evaluated for sensitivity, specificity, balanced accuracy, F1 score, and Youden’s J. The primary rule selected the threshold that maximized Youden’s J among candidates satisfying a minimum validation specificity of 0.50. This specificity value was used as a minimal guardrail against all-positive decisions in a weak-label setting, rather than as a clinically optimized deployment requirement. If no candidate satisfied the specificity constraint, the unconstrained Youden threshold was used. A fixed threshold of 0.5 and other validation-derived thresholds were retained as diagnostic comparisons, but the primary results use the specificity-constrained validation threshold. This threshold policy is part of the transfer protocol because weak-label adaptation can produce useful ranking while still collapsing to a single predicted class at an unsuitable operating point. Reporting AUC alone would therefore be insufficient; all thresholded metrics are tied to a stated threshold-selection rule.

### 4.7. Evaluation protocol

Animal-level partitioning. All partitions were defined at the cow level. Sequences from a given cow appeared in exactly one of training, validation, or held-out testing within a fold. The fixed held-out test set contained eight cows and was never used for training, model selection, calibration, or threshold selection. The remaining 23 cows formed the training pool. Five validation folds were constructed, each with four validation cows, so 20 training-pool cows served as validation cows once. The three remaining training-pool cows were included in the training subset of every fold and were never used for validation. This design preserves test-set independence and avoids repeated-identity leakage, while making explicit that validation coverage is not exhaustive over every non-test cow. It reduces identity, coat-pattern, background, and repeated-video leakage, a known concern in facial video analysis and external validation (Broome et al., 2023; Othman et al., 2019; Yu et al., 2022).

Aggregation levels. Predictions were evaluated at sequence, video, and cow levels. Sequence-level scores correspond to one 10 s face sequence. Because adjacent sequences extracted from the same video can share approximately 2 s of content, sequence-level results are treated as secondary descriptive evidence rather than as independent observations. Video- and cow-level scores were computed by averaging constituent sequence scores. Cow-level results are the primary animal-welfare reporting level because intervention decisions are made at the animal rather than frame level.

Metrics. Ranking was measured with the area under the receiver operating characteristic curve (AUC), with cow-level AUC treated as the primary animal-level ranking summary and sequence-level AUC reported as a secondary descriptor of score behavior. Thresholded performance was summarized by balanced accuracy, precision, recall, F1 score, specificity, and confusion counts. AUC and thresholded metrics answer different questions and are therefore reported together: AUC measures ranking ability, whereas thresholded metrics describe the operating behavior of the selected decision rule.

### 4.8. Statistical comparison and uncertainty analysis

Statistical analysis was designed around the dependency structure of the test set. Although the held-out set contains 143 sequences, these sequences are clustered within eight cows and include overlapping windows from the same source videos; treating sequences as independent would therefore overstate the effective sample size. The primary uncertainty analysis used cow-clustered bootstrap resampling. In each bootstrap replicate, held-out cows were sampled with replacement, and all sequences from the sampled cows were retained. Sequence-level metrics were recomputed from the pooled resampled sequences only as secondary descriptive summaries, and cow-level metrics were recomputed after averaging scores within each resampled cow. Percentile 95% confidence intervals were reported for the principal AUC estimates and for thresholded operating metrics where applicable.

Protocol comparisons used paired tests because all candidate protocols were evaluated on the same held-out cows and the same held-out sequences. For AUC differences, the primary test was a cow-clustered paired permutation procedure: for each cow, the two protocols’ score vectors were either kept in their original assignment or swapped as a block, and the difference in sequence AUC was recomputed. Because the test set contained eight cows, all cow-level swap patterns could be enumerated exactly. This procedure tests whether the observed AUC difference is larger than expected under exchangeability of protocol scores within the same animals, while preserving sequence clustering within cow.

Standard sequence-level AUC comparison tests were treated as sensitivity analyses rather than primary evidence because they do not fully address the clustered structure of repeated sequences from the same animals. The main claims therefore rely on cow-clustered confidence intervals and paired cow-clustered comparisons. When intervals are wide or paired comparisons do not support separation between protocols, the results are interpreted as differences in point estimates rather than statistically established superiority. Condition-specific summaries are reported descriptively because some health-condition strata contain few sequences and are not powered for independent inference.

## 5. Results

Results are reported on the fixed eight-cow held-out test set comprising 143 sequences, including 98 healthy-status proxy and 45 unhealthy-status proxy sequences. Evaluation was performed at both sequence and cow levels, with cow-level performance serving as the primary animal-level endpoint. Sequence-level AUC is reported as a secondary descriptor because adjacent 10 s windows extracted with an 8 s stride share approximately 2 s of content and are clustered within cows. All target-domain values measure discrimination of weak health-status proxies for pain-related discomfort, not validated pain detection. All configurations were assessed under identical animal-level partitions and validation procedures, with operating points set by the validation-selected threshold described in Section 4.6, enabling direct comparison of transfer protocols under a common evaluation framework (Figure 3).

**Figure 3.**
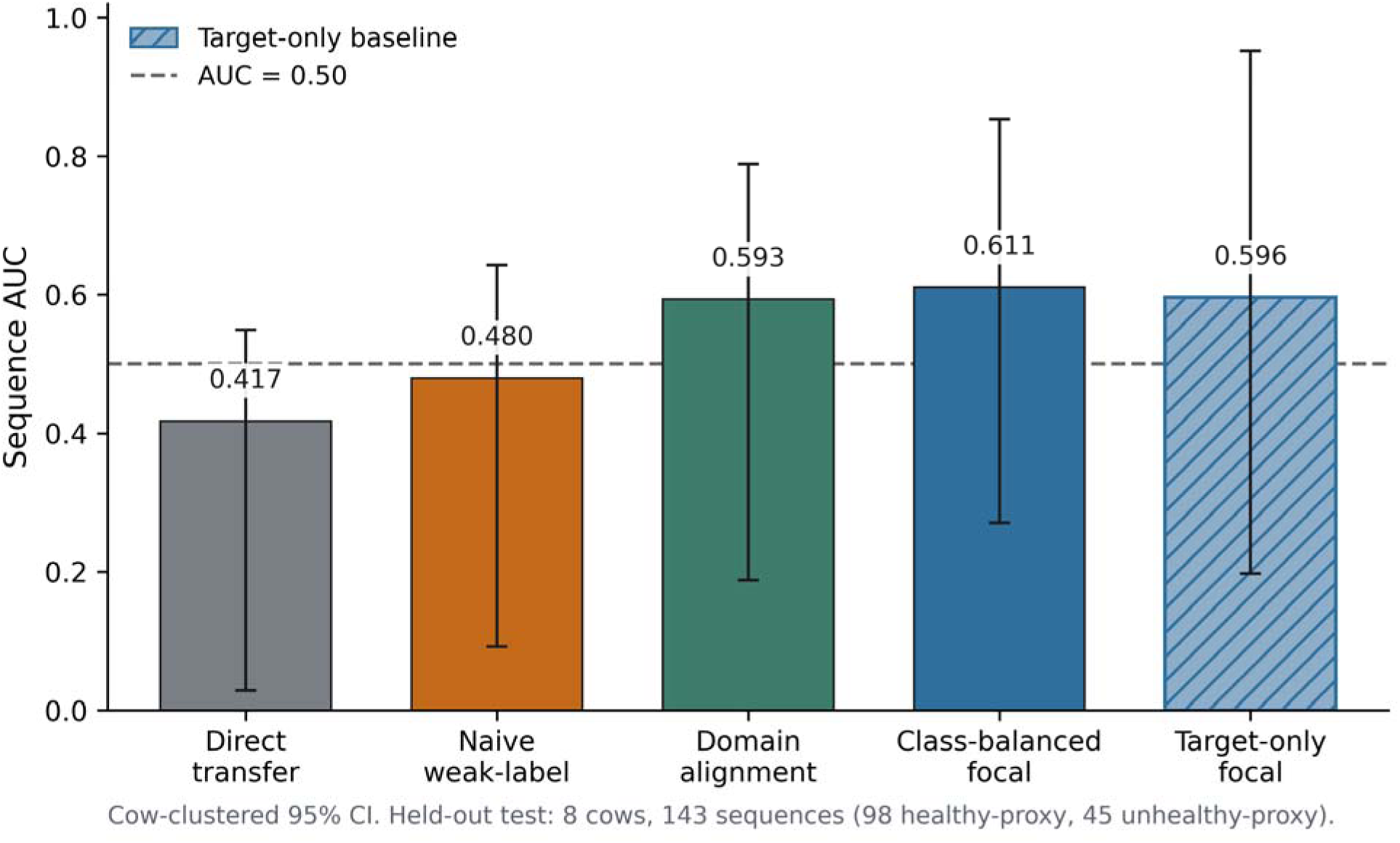
Performance of representative transfer protocols on the held-out dairy-cow test cohort. Sequence-level AUC for direct transfer, weak-label adaptation, domain-alignment, and class-balanced adaptation strategies evaluated on the same eight-cow held-out test set. Error bars indicate cow-clustered 95% confidence intervals, and the dashed line denotes chance performance (AUC = 0.50).

The results are organized around four findings. First, direct source-domain transfer was weak, indicating that the postoperative beef-cattle representation did not directly generalize to weakly labeled dairy-cow health states. Second, adaptation behavior was highly protocol dependent: naive weak-label objectives frequently collapsed to a single prediction class, while class-balanced focal objectives produced the most usable operating points. Third, ranking performance and decision behavior diverged; calibration preserved AUC but degraded thresholded predictions. Fourth, the target-only comparison showed that objective design and operating-point choices contributed more to observed performance than UCAPS source initialization under these weak-label conditions. These findings frame transferability not as a property that can be assumed from pretraining, but as an empirical claim that must be tested under animal-level uncertainty.

### 5.1. Limited generalization of source-domain representations

The unadapted source representation generalized weakly to dairy health-status proxies. Direct transfer reached sequence AUC 0.418 [0.028, 0.550] and cow-level AUC 0.400 [0.000, 0.857]. Mean source-model scores were tightly concentrated across the test cows, indicating weak separation of healthy-proxy and unhealthy-proxy sequences. Thus, the facial pain representation learned from controlled postoperative beef cattle did not, on its own, rank dairy-cow discomfort proxies above chance under simultaneous breed, sex, environment and label shift. This source-only result establishes the baseline against which all adaptation protocols should be interpreted. It also motivates the target-only comparison in Section 5.8, which tests whether UCAPS initialization contributes more than objective design and threshold policy when the target labels are weak.

### 5.2. Weak supervision alone is insufficient for reliable transfer

Weak-label adaptation with standard objectives and no class balancing produced near-chance ranking and degenerate decision behavior, revealing the first failure mode, threshold collapse, in which the adapted model defaults to a single prediction class. Binary cross-entropy reached sequence AUC 0.480 [0.092, 0.643], generalized cross-entropy reached 0.477 [0.089, 0.639], and focal loss reached 0.487 [0.127, 0.661]. The first two configurations predicted no positive sequences at the validation-selected threshold, yielding zero true positives among 45 unhealthy-status proxy sequences. The focal-loss configuration produced only two true positives. Cow-level AUC was 0.400 for binary cross-entropy and generalized cross-entropy and 0.333 for focal loss. These configurations illustrate why ranking metrics alone are insufficient in weak-label livestock welfare tasks. Although their sequence-level AUCs were close to chance rather than uniformly zero, their thresholded behavior was operationally unusable because all or nearly all sequences were assigned to one class. Weak supervision can therefore preserve a source representation while still failing as a screening decision rule.

### 5.3. Domain alignment partially mitigates distribution shift

A source-target alignment term improved sequence-level ranking relative to naive weak-label adaptation but did not yield a robust operating point. Correlation alignment reached sequence AUC 0.593 [0.188, 0.789] in the representative setting, and adversarial alignment ranged from 0.591 to 0.593. However, the operating behavior remained unstable. The best adversarial setting produced F1 0.361, recall 0.333, and 15 true positives, whereas the correlation-alignment settings remained near-degenerate, with F1 0.136, recall 0.089, and four true positives. Cow-level AUC was 0.400 for these alignment configurations. A separate alignment evaluation on the same held-out cows regressed to sequence AUC 0.467-0.470 and all-negative collapse. Thus, alignment improved ranking in selected configurations but did not consistently prevent collapse, and it did not improve cow-level discrimination. This result indicates that feature alignment alone is insufficient when source and target labels differ in biological meaning and fidelity.

### 5.4. Objective design strongly influences transfer performance

Among the evaluated adaptation strategies, objective design emerged as the strongest determinant of transfer behavior. The class-balanced focal configuration with gamma = 2.5 achieved the highest sequence-level discrimination and produced the most stable operating point on the held-out dairy-cow cohort. Sequence-level AUC reached 0.611 [0.271, 0.853], while cow-level AUC reached 0.667 [0.200, 1.000]. Unlike the naive objectives and several alignment configurations, this protocol maintained meaningful sensitivity and avoided collapse toward a single prediction class.

At the sequence level, this configuration reached balanced accuracy 0.593, F1 0.466, recall 0.533, and precision 0.414, with a confusion count of 24 true positives, 21 false negatives, 34 false positives, and 64 true negatives. At the cow level, it reached balanced accuracy 0.800, F1 0.750, and recall 1.000. The confusion-count behavior of this and the other protocols at the validation-selected threshold is shown in Figure 4. In a paired cow-clustered comparison against the representative alignment result, the cow-level AUC difference was small and not statistically reliable (delta = 0.017; p = 0.977), confirming that even among transfer protocols, apparent differences were small relative to uncertainty.

**Figure 4.**
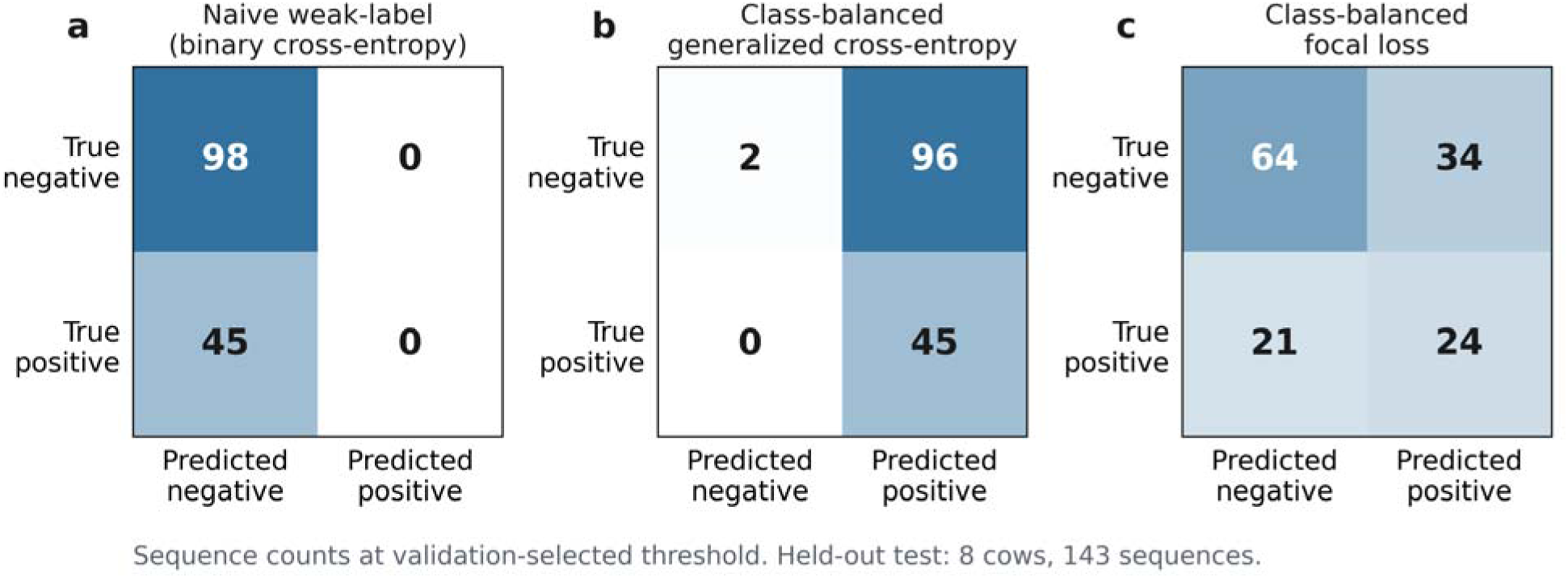
Representative operating-point behaviors under different adaptation objectives. Confusion matrices for naïve weak-label adaptation, class-balanced generalized cross-entropy, and class-balanced focal adaptation on the held-out dairy-cow cohort. The examples illustrate distinct operating regimes, including all-negative collapse, all-positive collapse, and stable class discrimination.

Objective design also separated usable from degenerate behavior. A class-balanced generalized cross-entropy objective with q = 0.6 produced all-positive collapse, with recall 1.000 but 96 false positives and only two true negatives. A lower focal focusing parameter reached sequence AUC 0.543. These contrasts show that class balancing alone was insufficient; the specific class-balanced focal objective produced the strongest and most stable transfer behavior. The main descriptive results for all protocol families on the held-out test set are reported in Table 5.

**Table 5.** Comparison of protocol-driven transfer strategies under identical animal-level evaluation conditions on the eight-cow held-out test set, isolating how each adaptation choice governs transfer.

| Configuration family | Representative configuration | Sequence AUC [95% CI] | Sequence F1 | Sequence recall | Cow AUC [95% CI] |
| --- | --- | --- | --- | --- | --- |
| Direct transfer | Source model, no target update | 0.418<br>[0.028, 0.550] | not applicable | not applicable | 0.400<br>[0.000, 0.857] |
| Naive weak-label adaptation | Binary cross-entropy | 0.480<br>[0.092, 0.643] | 0.000 | 0.000 | 0.400<br>[0.000, 0.750] |
| Naive weak-label adaptation | Generalized cross-entropy | 0.477<br>[0.089, 0.639] | 0.000 | 0.000 | 0.400<br>[0.000, 0.750] |
| Naive weak-label adaptation | Focal loss | 0.487<br>[0.127, 0.661] | 0.074 | 0.044 | 0.333<br>[0.000, 0.750] |
| Domain alignment | Correlation alignment | 0.593<br>[0.188, 0.789] | 0.136 | 0.089 | 0.400<br>[0.000, 0.938] |
| Domain alignment | Adversarial alignment | 0.593<br>[0.174, 0.795] | 0.361 | 0.333 | 0.400<br>[0.000, 0.833] |
| Objective design | Class-balanced focal, gamma = 1.5 | 0.543<br>[0.300, 0.852] | 0.385 | 0.444 | 0.667<br>[0.333, 1.000] |
| Objective design | Class-balanced focal, gamma = 2.5 | 0.611<br>[0.271, 0.853] | 0.466 | 0.533 | 0.667<br>[0.200, 1.000] |

Across transfer families, class-balanced focal objectives produced the only consistently non-degenerate operating regimes. However, even the best transfer configuration reached cow-level AUC 0.667 with wide confidence intervals, and its advantage over alternative transfer protocols was not statistically reliable. The target-only comparison in Section 5.8 further shows that this usable behavior was driven more by objective and protocol design than by source initialization.

### 5.5. Calibration leaves ranking intact but can break the operating point

Post hoc temperature scaling demonstrated that discrimination and decision quality can diverge. For the class-balanced focal configuration, AUC remained 0.611 after calibration, but F1 dropped from 0.466 to 0.038 and recall dropped from 0.533 to 0.022, corresponding to one true positive. This is the second failure mode, calibration-induced collapse, in which score ranking is preserved but thresholded decision behavior deteriorates. Several alignment configurations also shifted to all-positive decisions after calibration. This result is important because livestock monitoring systems act on decisions, not only on ranks. A model can therefore appear unchanged under AUC while becoming unusable at the selected operating point. Because calibration changed thresholded behavior without improving ranking, raw operating-point metrics are used for the primary conclusions, while calibrated metrics are treated as diagnostics of score instability under weak labels. The effect of temperature scaling on the score distributions and the selected operating point is shown in Figure 5.

**Figure 5.**
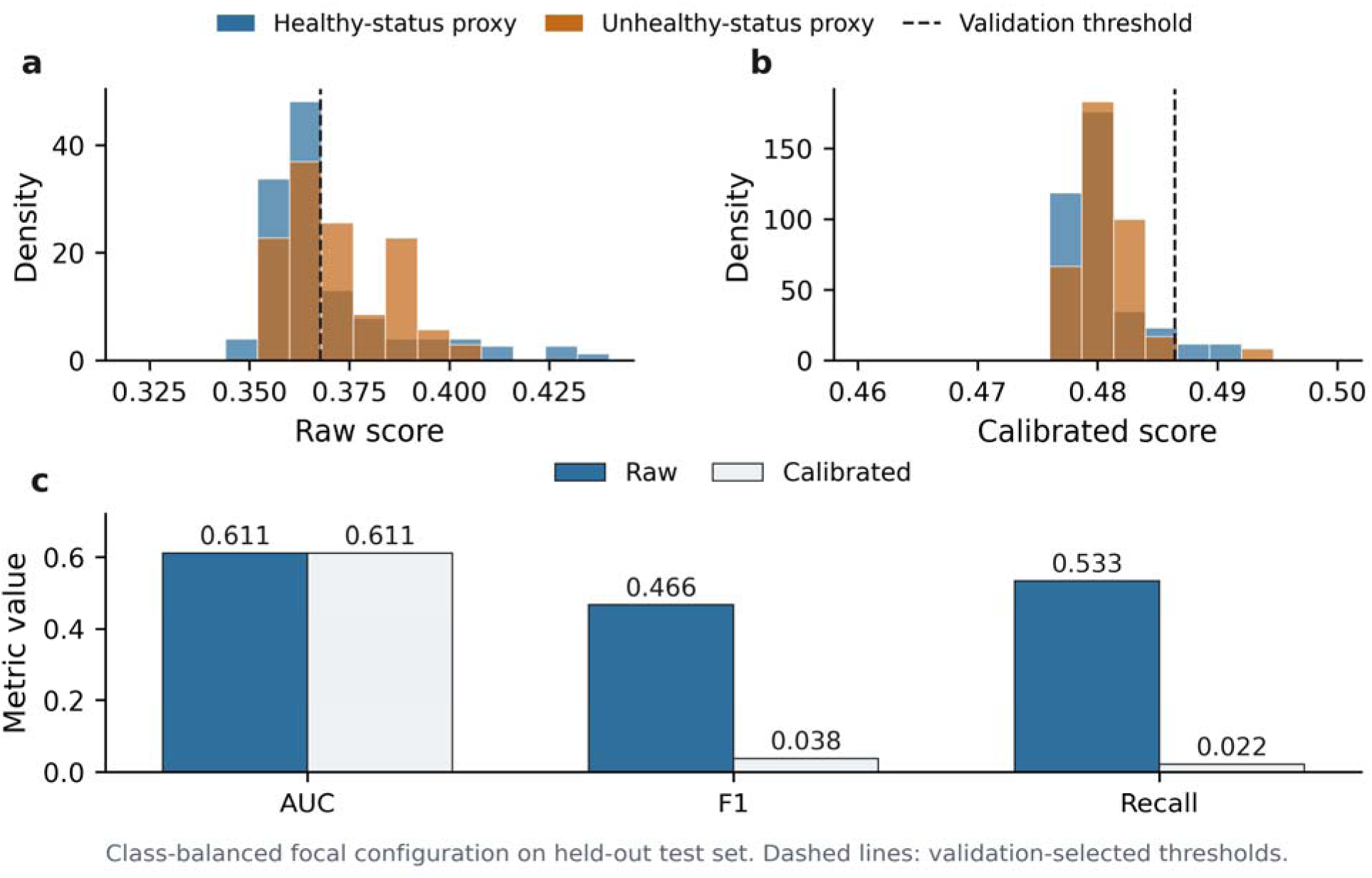
Influence of calibration on decision behavior. Raw and temperature-scaled prediction-score distributions for the class-balanced focal configuration. Although calibration preserves ranking performance, it substantially alters thresholded decision behavior on the held-out test set.

### 5.6. Transferability varies across health condition

Stratifying the class-balanced focal predictions by health condition showed an asymmetry across target conditions. Possible mastitis sequences were flagged positive at a higher rate (0.586; n = 29) than lameness sequences (0.438; n = 16), while healthy-status proxy sequences were flagged at 0.345 (n = 87). The full stratified rates are reported in Supplementary Table S1 and the operating behavior is shown in Supplementary Figure S2. The strata are small and are therefore reported as condition-level flagging rates rather than powered comparisons. Although underpowered, this pattern is biologically informative. Mastitis is a systemic inflammatory condition that may influence facial expression, head posture and general demeanor, whereas lameness often presents more strongly through gait, posture and weight-bearing behavior. The observed asymmetry therefore suggests that facial-only models may transfer more readily for conditions with systemic or affective facial manifestations than for locomotion-dominated disorders.

### 5.7. Animal-level uncertainty remains substantial

Animal-level uncertainty was large relative to the observed effects. For class-balanced focal adaptation, cow-level AUC was 0.667 with a bootstrap 95% confidence interval of [0.20, 1.00] (Figure 6), spanning from below chance to perfect ranking. The point estimate sits 0.167 above chance, while the interval extends roughly 0.47 below and 0.33 above it. Direct transfer and the representative alignment configuration each had cow-level AUC 0.400 with intervals spanning chance and near-complete uncertainty. Paired cow-clustered comparisons did not establish statistically reliable superiority for any observed AUC differences, including the comparison between the best alignment result and class-balanced focal adaptation. Inner-validation AUC also varied widely across folds, and high validation AUC did not consistently translate to the held-out cows. In other words, the animal-level uncertainty band was wider than the apparent transfer gain. These findings show why strict animal-held-out validation is essential: sequence-level improvements can suggest progress while individual-animal evidence remains underpowered.

**Figure 6.**
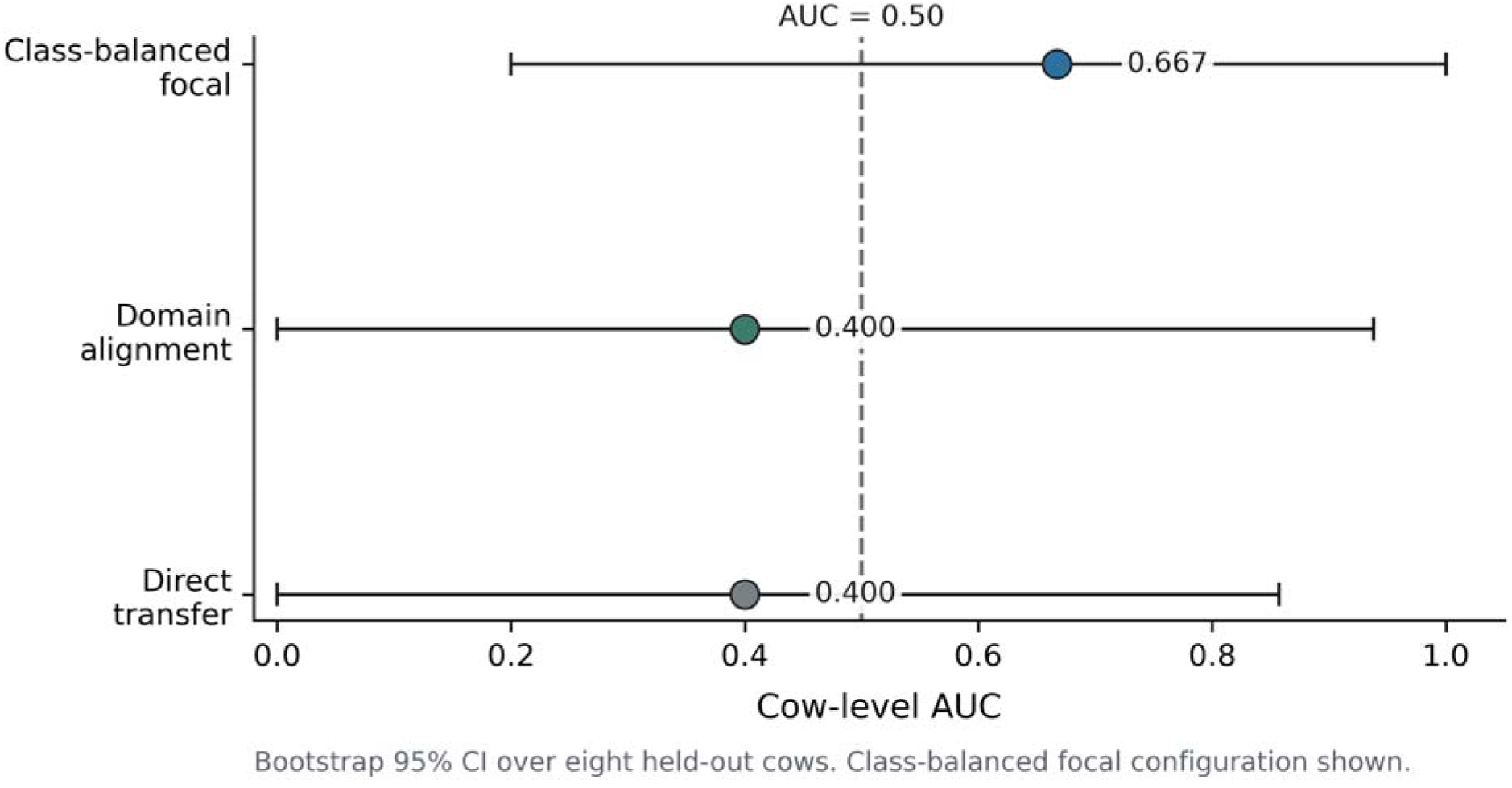
Animal-level uncertainty across protocol configurations. Cow-level AUC estimates with bootstrap 95% confidence intervals over the eight held-out cows for pain-related discomfort proxies. Wide intervals indicate that animal-level uncertainty exceeds the observed differences among protocol families.

### 5.8. Comparison with target-only dairy learning

To determine whether UCAPS pretraining contributed more than protocol design, we trained the same fixed architecture from random initialization on the identical 549-sequence material and eight-cow held-out test set, using only weak video_health_status labels and no UCAPS checkpoint. This target-only baseline used the same cow-held-out split, class-balanced sampling, and validation-threshold policy as the transfer experiments, but omitted source initialization and did not freeze the convolutional encoder, because freezing is inappropriate when weights are untrained. Three objective variants were evaluated: binary cross-entropy, class-balanced focal loss with focusing parameter gamma = 2.5, matching the best transfer protocol except for initialization, and class-balanced generalized cross-entropy with q = 0.6. The target-only results on the held-out test set are reported in Table 6.

**Table 6.** Contribution of source pretraining isolated: target-only (from-scratch) weak-label results under the same animal-level evaluation as the transfer protocols.

| Configuration | Initialization | Sequence AUC [95% CI] | Sequence F1 | Sequence Recall | Cow AUC [95% |
| --- | --- | --- | --- | --- | --- |

|  |  |  |  |  | **CI]** |
| --- | --- | --- | --- | --- | --- |
| Target-only<br>weak-label | Random / from-scratch,<br>BCE + class balancing | 0.502 [0.152, 0.967] | 0.189 | 0.111 | 0.667<br>[0.143,<br>1.000] |
| Target-only<br>weak-label | Random / from-scratch,<br>focal $\gamma = 2.5$ + class<br>balancing | 0.596 [0.197, 0.953] | 0.532 | 0.733 | 0.667<br>[0.143,<br>1.000] |
| Target-only<br>weak-label | Random / from-scratch,<br>GCE $q = 0.6$ + class<br>balancing | 0.487 [0.162, 0.863] | 0.071 | 0.044 | 0.667<br>[0.188,<br>1.000] |

The results show that, under the same protocol logic, target-only training matched the best transfer configuration at the cow level and nearly matched it at the sequence level. This comparison is central to the interpretation of the study because it separates the effect of source initialization from the effect of objective design and operating-point selection.

Target-only class-balanced focal training produced a non-degenerate operating point without UCAPS initialization, with F1 0.532, recall 0.733, and confusion count 33/12/46/52. This mirrors the objective sensitivity observed in Sections 5.2-5.4: naive or mismatched objectives collapsed or remained near chance, whereas class-balanced focal produced the most usable behavior. At the cow level, the best target-only configuration matched the best transfer configuration at AUC 0.667, with comparably wide bootstrap intervals [0.143, 1.000] versus [0.200, 1.000] for UCAPS-initialized class-balanced focal adaptation.

In the direct comparison between target-only and transfer class-balanced focal adaptation, sequence AUC was 0.596 for target-only focal versus 0.611 for transfer focal, a difference of 0.015 that was not statistically reliable under the cow-clustered paired permutation test (p = 0.984). Taken together, these comparisons show that, in this setting, objective design and evaluation protocol exerted a larger influence on ranking and operating-point behavior than UCAPS initialization itself, while animal-level uncertainty remained substantial in both cases.

## 6. Discussion

The central contribution of this study is PDTE, a protocol-driven framework for determining when transfer claims in livestock welfare AI are genuine, weak, or artefacts of adaptation choices. Applied to bovine facial pain transfer from postoperative beef cattle to weakly labeled dairy farm video, PDTE’s most important empirical findings are not a headline sequence AUC of 0.611 but the failure modes and uncertainty structure that govern whether a system would behave sensibly in a farm-monitoring context. Naive weak-label adaptation collapsed to threshold degeneracy; temperature scaling produced calibration-induced collapse in which ranking was preserved yet operating behavior was destroyed; performance was highly sensitive to objective and threshold choices; and cow-level bootstrap intervals (0.20-1.00) were wider than the differences among protocol families. Respectable sequence-level AUC can therefore coexist with operationally unusable decisions when threshold selection and calibration are not evaluated jointly with discrimination a conclusion that would be missed if transfer were reported as a single end-to-end score. We organize the discussion around PDTE as the methodological contribution, the bovine case study as the evidence base, and the implications for agricultural domain shift, deployability assessment, and multimodal welfare sensing.

### 6.1. Protocol-guided evaluation of transferability in livestock welfare AI

The primary contribution is methodological, and the bovine transfer experiment is its test case. Where prior livestock computer-vision studies report a single end-to-end performance number, PDTE decomposes transfer into label mapping, objective, alignment, model selection, calibration, and threshold policy, and evaluates each component under identical animal-level external validation. This decomposition is what makes the results interpretable: the same dataset and architecture yield collapse, partial recovery, or a usable operating point depending entirely on which component is varied, so apparent transfer success or failure is shown to be a property of the protocol rather than of the representation or source pretraining alone. In this cohort, objective design was the dominant lever, class-balanced focal adaptation reached the best operating point (sequence AUC 0.611, cow-level AUC 0.667), and a from-scratch target-only model essentially matched it (sequence AUC 0.596; paired p = 0.984), indicating that the added value of UCAPS source pretraining is limited relative to careful protocol design under weak agricultural labels. PDTE is reusable beyond this case: any agricultural transfer setting can be analyzed by holding the model fixed and treating these components as the experimental variables.

These findings are particularly relevant for precision livestock farming, where models developed under controlled experimental conditions are increasingly expected to operate across farms, breeds, management systems, and welfare contexts. The ability to evaluate transferability systematically may therefore be as important as the development of increasingly complex model architectures.

### 6.2. Characterizing agricultural domain shift

The results characterize the domain gap that agricultural welfare AI must cross, and show that it is partly covariate and partly conceptual. Domain alignment, which targets appearance and acquisition differences (Ganin et al., 2016; Ben-David et al., 2010), raised sequence-level ranking above naive weak-label adaptation, indicating that some transferable structure remains in the source representation. Yet alignment did not produce a dependable operating point, consistent with theory when part of the discrepancy is conceptual: the source labels encode a staged postoperative pain trajectory in male beef bulls, whereas the target labels encode heterogeneous disease-associated states in female dairy cows whose facial correlates need not match the source construct (Mogil et al., 2020; Mota-Rojas et al., 2025). The bull-to-cow contrast is an additional uncontrolled axis alongside breed, etiology, and environment. The transferable component therefore appears to be correlate structure rather than label equivalence: the source representation was reused, but not reproduced as a validated target-domain pain detector. This simultaneous multi-axis shift is the regime most agricultural deployments actually face, and PDTE makes its effect measurable rather than implicit.

### 6.3. Transfer failure modes as design principles

PDTE exposed two characteristic failure modes in weak-label livestock welfare AI, each of which yields a concrete design principle. Threshold collapse, in which naive weak-label adaptation defaults to a single prediction class, arises when healthy-proxy and unhealthy-proxy facial score distributions overlap heavily under weak labels; the design response is to treat class-balanced objectives and operating-point diagnostics as defaults rather than refinements. Calibration-induced collapse, in which temperature scaling preserves AUC yet compresses probabilities so that the selected operating point loses recall, shows that better-looking probabilities are not better decisions; the design response is to evaluate ranking, calibration, and thresholded behavior as separate quantities. These failure modes are consistent with the challenges expected under weak agricultural labels and small validation sets and naming them gives future systems a checklist of what to test before deployment.

### 6.4. Animal-level uncertainty-aware validation

The study makes a case that animal-level, uncertainty-aware evaluation should be a standard rather than an option for livestock welfare AI. Cow-clustered bootstrap intervals were wide (cow-level AUC 0.667 [0.20, 1.00]), validation AUC varied across folds, and high validation AUC did not consistently translate to held-out cows, so a sequence-level point estimate read without animal-level uncertainty would substantially overstate what the model can do for an individual cow. This reinforces, in a cross-domain bovine setting, the subject-level evaluation emphasized across animal and human pain recognition (Broome et al., 2023), and the cross-database degradation documented in human facial-video pain analysis (Othman et al., 2019) and in medical-imaging external validation (Yu et al., 2022). Relative to in-domain cattle pain studies reporting stronger controlled-condition performance (Feighelstein et al., 2026), the contribution here is to quantify what survives transfer and to bound it honestly with animal-level uncertainty.

### 6.5. Transferability across conditions and multimodal sensing

Transferability varied across health conditions. The class-balanced focal model flagged possible mastitis sequences more often than lameness sequences, consistent with prior evidence that mastitis-related discomfort alters dairy-cow facial expression (Ginger et al., 2023) whereas lameness is expressed more through gait and posture (Flower and Weary, 2008). The conserved part of the facial signal therefore appears to be whatever resembles acute, systemically expressed pain, while posture-dominated conditions are poorly captured by the face alone. This motivates multimodal welfare sensing in which facial video is one channel among locomotion, feeding and rumination behavior, posture, thermal imaging, and clinical records (Rohan et al., 2024; Afonso et al., 2020), so that conditions poorly expressed in the face are still detected.

### 6.6. Toward a deployability-assessment framework for agricultural AI

Taken together, these contributions sketch a deployability-assessment framework: before a livestock welfare model is deployed, transfer should be decomposed into its protocol components, stress-tested for threshold and calibration collapse, and reported with animal-level uncertainty under external validation. The framework also reframes a common objection, that one should simply train a dairy-specific model rather than transfer from beef cattle. Because commercial dairy operations rarely hold hundreds of time-aligned validated pain annotations, the realistic regime is weak, video-level health-status proxies; under naive objectives this regime collapsed whether training started from a UCAPS checkpoint or from random initialization. The target-only baseline (Section 5.8) matched the transfer model at the animal level (cow-level AUC 0.667; sequence AUC 0.596 versus 0.611; p = 0.984), so under these conditions adaptation protocol, not source pretraining, governed usable behavior. Source pretraining may help in other cohorts or label regimes, but these findings underscore that farm-scale deployment will require larger multi-herd validation and likely multimodal integration. The practical lesson for agricultural AI is that protocol design and honest animal-level evaluation, captured by PDTE, defines a validation path by which future welfare models can be evaluated for deployment.

## 7. Limitations

The scope of this demonstration defines clear boundaries on its claims, and each points to a concrete extension of PDTE. The target labels are weak, video-level health-status proxies rather than time-aligned pain scores; the framework treats this transparently by reporting proxy discrimination, and pairing the same protocol with independently adjudicated labels would separate genuine transfer failure from label noise. The held-out cohort comprises eight cows, which is why the cow-level intervals are wide and no protocol is declared superior; the cow-clustered bootstrap and paired animal-level tests make this uncertainty explicit and should narrow as PDTE is applied to larger multi-herd cohorts. Because the 10 s windows were extracted with an 8 s stride, adjacent sequences are not independent, which is why animal-level metrics are primary and a non-overlapping-window analysis is a natural sensitivity extension. By design the study fixes one source representation, one architecture, and a prespecified selection rule so that the protocol is the only experimental variable; the same framework can now evaluate alternative backbones, pretraining strategies, and selection schemes as added factors. Full external replication depends on access to the source model and protected farm video, but the protocol, splits, and evaluation rules are fully specified, so the methodology is portable even where these data cannot be shared.

## 8. Conclusions

Transferability in livestock welfare AI cannot be inferred from source-domain performance, model architecture, or the availability of a pretrained representation. It must be demonstrated under realistic biological, environmental, and label-quality shifts, with evaluation performed at the animal level and with explicit uncertainty. The Protocol-Driven Transfer Evaluation (PDTE) framework provides such a methodology by decomposing adaptation into auditable protocol components, including label mapping, objective design, domain alignment, model selection, calibration, and threshold policy, and evaluating each under identical animal-level external validation.

Applied to bovine facial pain transfer from postoperative beef cattle to weakly labeled dairy-farm video, PDTE showed that the principal empirical contribution is not a headline sequence AUC, but the characterization of transfer limits, decision failure modes, and animal-level uncertainty. Direct source-only transfer was weak, with sequence AUC 0.418 and cow-level AUC 0.400. Class-balanced focal adaptation produced the strongest transfer operating point, reaching sequence AUC 0.611 and cow-level AUC 0.667. However, target-only focal training achieved comparable performance without source initialization, with sequence AUC 0.596 and no reliable paired difference from transfer adaptation (p = 0.984). Under these weak-label conditions, protocol design and operating-point management therefore contributed more to usable behavior than source-domain pretraining.

PDTE also documented two important failure modes in weak-label agricultural transfer: threshold collapse, in which adaptation defaults to a single prediction class, and calibration-induced collapse, in which score ranking is preserved while decision behavior deteriorates. These results show that ranking metrics alone are insufficient for livestock welfare screening, where thresholded decisions determine whether animals are flagged for attention.

Animal-level uncertainty remained substantial, with the best cow-level AUC confidence interval spanning 0.20 to 1.00. This uncertainty limits deployment claims and reinforces the need for larger multi-herd validation, stronger clinical labels, and likely multimodal integration before facial-video systems are considered for farm-scale welfare monitoring. Within these boundaries, PDTE offers a reusable framework for determining when agricultural welfare-AI transfer claims are genuine, fragile, or protocol-driven before such systems reach on-farm use.

## Data and model availability

The source UCAPS-associated cattle pain dataset and the target RAC dairy-cow video dataset are not publicly deposited because they contain animal-health information, facility-specific visual material, and data governed by study permissions, ethics approvals, and institutional data-use conditions. Requests for access to these datasets may be directed to the corresponding author and will be considered on a case-by-case basis, subject to the relevant ethical, legal, institutional, and data-sharing requirements. To support verification and reproducibility, de-identified derived materials underlying the analyses, including animal-level split definitions, protocol configuration files, model prediction tables, threshold-selection outputs, and statistical-analysis results, will be made available upon reasonable request where permitted. Source model weights and target-derived face-sequence data may also be shared with qualified researchers under an approved data-use agreement, provided that sharing is consistent with the applicable ethics approvals, facility permissions, and animal-health data restrictions.

## Acknowledgements

The authors sincerely thank the Natural Sciences and Engineering Research Council of Canada and the Nova Scotia Department of Agriculture for funding this study.

## CRediT authorship contribution statement

Shivam Patel: Data curation, Formal analysis, Investigation, Methodology, Validation, Visualization, Writing – original draft. Suresh Neethirajan: Conceptualization, Funding acquisition, Project administration, Resources, Supervision, Writing – review & editing.

