## Supplementary Material for "Protocol-Guided Cross-Domain Transfer Learning for Bovine Facial Pain Recognition under Weak Dairy-Farm Labels"

### Supplementary Methods

#### S1. Transfer-protocol formalization

Each transfer system is formally described by the protocol tuple P = (M, L, A, S, C, T), applied to a fixed source-initialized model and fixed animal-level partitions. M is the target-label mapping, L the target objective, A an optional source-target alignment term, S the validation-based model-selection functional, C an optional calibration rule, and T the operating-threshold policy. Two systems with the same architecture and data partitions but a different value of any component are treated as different transfer protocols, so each main-text result traces to a defined substitution rather than to an unstructured set of implementation changes.

Model selection used validation animals only. The selection functional was S = 0.45 x AUC_sequence + 0.35 x AUC_cow + 0.20 x balanced accuracy, computed on validation animals. The weights are heuristic, were not tuned on the held-out test cows, and prioritize cow-level ranking (the welfare-relevant unit) while penalizing collapsed operating points through the balanced-accuracy term.

### Table S1. Exploratory condition-stratified positive prediction rate under class-balanced focal adaptation

| Group | n sequences | Fraction predicted positive |
| --- | --- | --- |
| Possible mastitis | 29 | 0.586 |
| Lameness | 16 | 0.438 |
| Healthy-status proxy | 87 | 0.345 |

Fraction of sequences predicted positive under class-balanced focal adaptation (gamma = 2.5) on the eight-cow held-out test set, stratified by health-condition group. The analysis is descriptive only and not powered for condition-level inference.

#### Table S2. Transfer-protocol components evaluated in this study

| Component | Definition | Values evaluated |
| --- | --- | --- |
| M: target-label mapping | Dairy health status to binary target label | Healthy-status to no-pain-related-discomfort proxy; unhealthy-status to pain-related-discomfort proxy |
| L: target objective | Loss on the target binary head | Binary cross-entropy, focal, or generalized cross-entropy, with or without class balancing |
| A: alignment term | Optional source-target feature-distribution penalty | None, adversarial domain alignment (DANN), or correlation alignment (CORAL) |
| S: selection functional | Validation criterion to choose a fold model | Weighted sequence AUC, cow AUC, and balanced accuracy (see formula above) |
| C: calibration rule | Optional score calibration fitted without test data | None or temperature scaling on validation logits |
| T: threshold policy | Continuous score to binary decision | Youden's J under validation specificity >= 0.50, with prespecified fallbacks |

#### Table S3. Model and adaptation hyperparameters

| Setting | Value |
| --- | --- |
| Input frames per sequence | 32 RGB frames, 112 x 112 px |
| Frame encoder embedding | 256-dimensional per frame (2D CNN, encoder frozen unless stated) |
| Temporal module | Single-layer recurrent unit, 128 hidden units, with attention pooling |
| Focal loss focusing parameter (gamma) | 1.5 and 2.5 |
| Generalized cross-entropy robustness (q) | 0.6 and 0.8 |
| Class-balanced effective-number weighting (beta) | 0.999 |
| Correlation alignment (CORAL) weight | 0.01, 0.02, 0.05, 0.10 |
| Adversarial alignment (DANN) domain weight | 0.10, 0.15, 0.20, 0.25, 0.50 |
| Calibration | Temperature scaling fitted on validation logits (monotonic; AUC unchanged) |
| Ensembling | Fold-level logits averaged before sigmoid |

All values are fixed study settings; the network topology and source initialization were held constant across every configuration, and only the protocol components in Supplementary Table S2 varied.

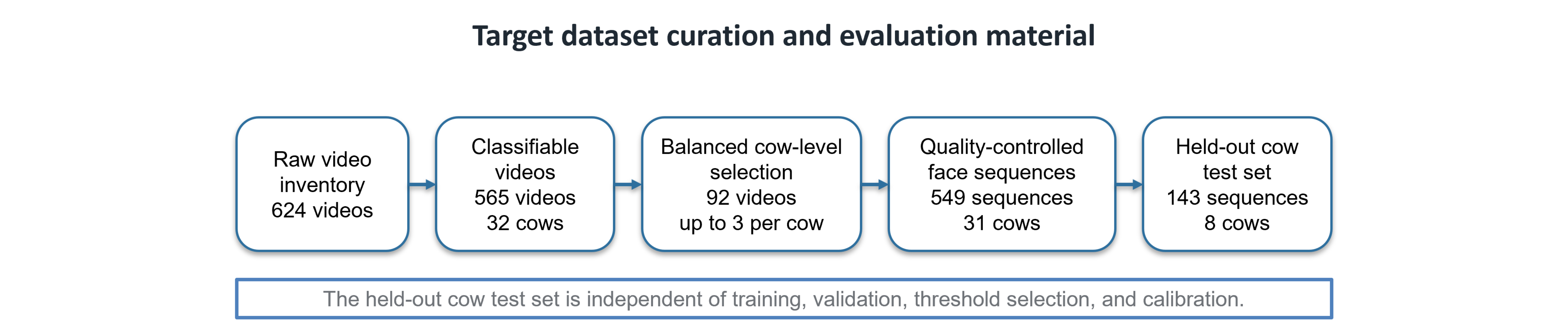

Figure S1. Target dataset curation and held-out evaluation material. Flow diagram summarizing the target dataset curation pathway from raw video inventory to classifiable videos, balanced cow-level selection, quality-controlled face sequences, and the held-out cow test set. The held-out cow test set was not used for training, validation, threshold selection, or calibration.

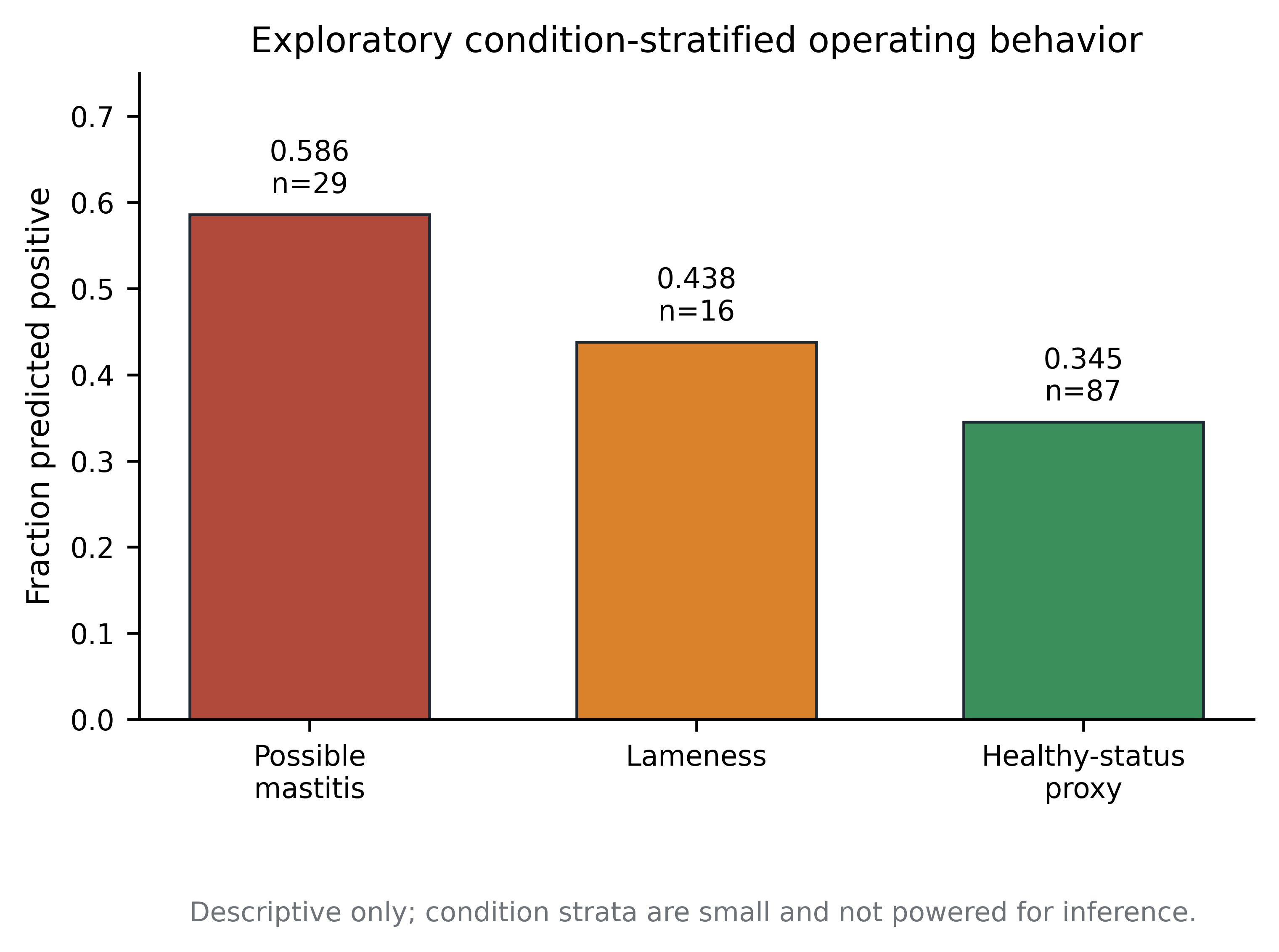

Figure S2. Exploratory condition-stratified operating behavior. Fraction of sequences predicted positive under the class-balanced focal configuration, stratified by selected health-condition groups. Possible mastitis sequences were flagged positive at a higher rate than lameness sequences and healthy-status proxy sequences. This analysis is descriptive only because the condition strata are small and not independently powered for inference.
